## Supplemental Figures for "Periplasm homeostatic regulation maintains spatial constraints essential for cell envelope processes and cell viability"

**Table S1 Proteomic results**

**Table S2. Substantive changes in steady-state protein levels in cell envelope of $Lpp^{+21}$**

**Table S3: Proteins used in the generation of the sequence similarity network**

**Table S4: Representative Lpp protein information**

**Table S5: Bacterial strains and primers used in the study**

**Table S6: Genes up-regulated in Lpp+21 strain**

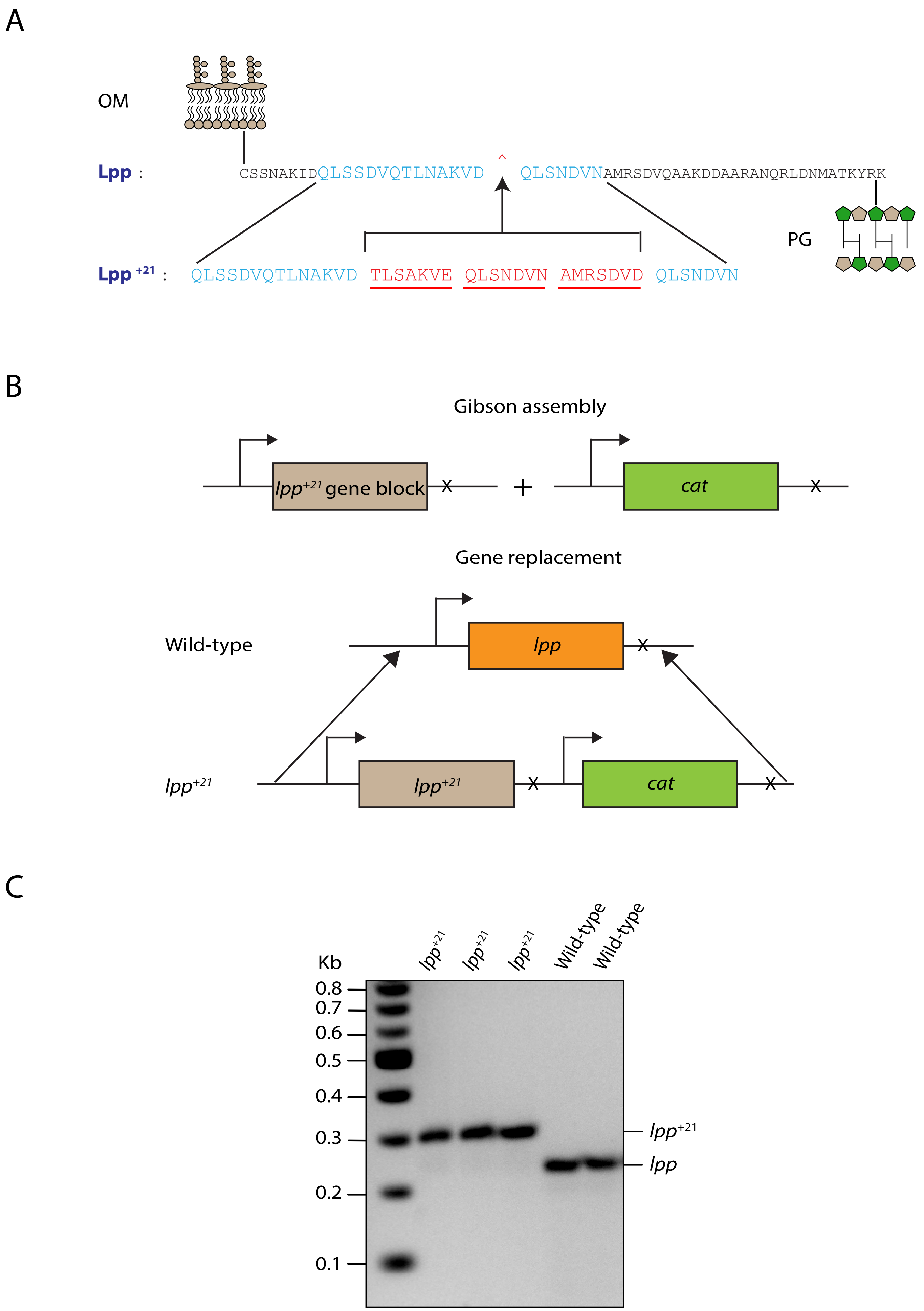
A
OM
Lpp :
CSSNAKIDQLSSDVQTLNAKVD ^ QLSNDVNAMRSDVQAAKDDAARANQRLDNMATKYRK
PG
Lpp +21 :
QLSSDVQTLNAKVD TLSAKVE QLSNDVN AMRSDVD QLSNDVN
B
Gibson assembly
lpp+21 gene block
cat
Gene replacement
Wild-type
lpp
lpp+21
lpp+21
cat
C
lpp+21
lpp+21
lpp+21
Wild-type
Wild-type
Kb
0.8
0.7
0.6
0.5
0.4
0.3
0.2
0.1
lpp+21
lpp

Figure S1

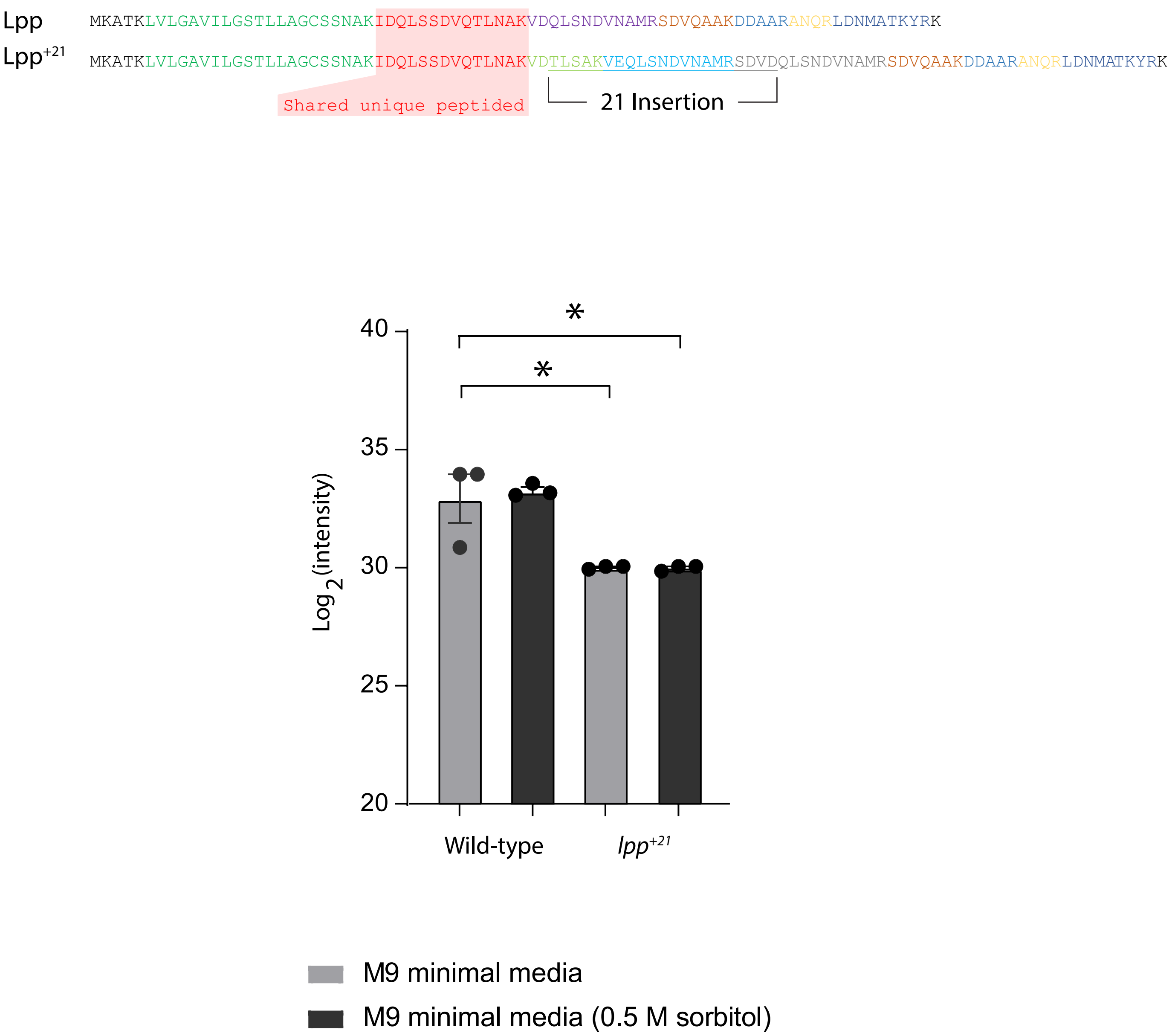
Lpp MKATKLVLGAVILGSTLLAGCSSNAKIDQLSSDVQTLNAKVDQLSNDVNAMRSDVQAAKDDAARANQRLDNMATKYRK
Lpp+21 MKATKLVLGAVILGSTLLAGCSSNAKIDQLSSDVQTLNAKVDTLSAKVEQLSNDVNAMRSDVDQLSNDVNAMRSDVQAAKDDAARANQRLDNMATKYRK
Shared unique peptided
21 Insertion
40
35
30
25
20
Log 2(intensity)
*
*
Wild-type
lpp+21
M9 minimal media
M9 minimal media (0.5 M sorbitol)

Figure S2

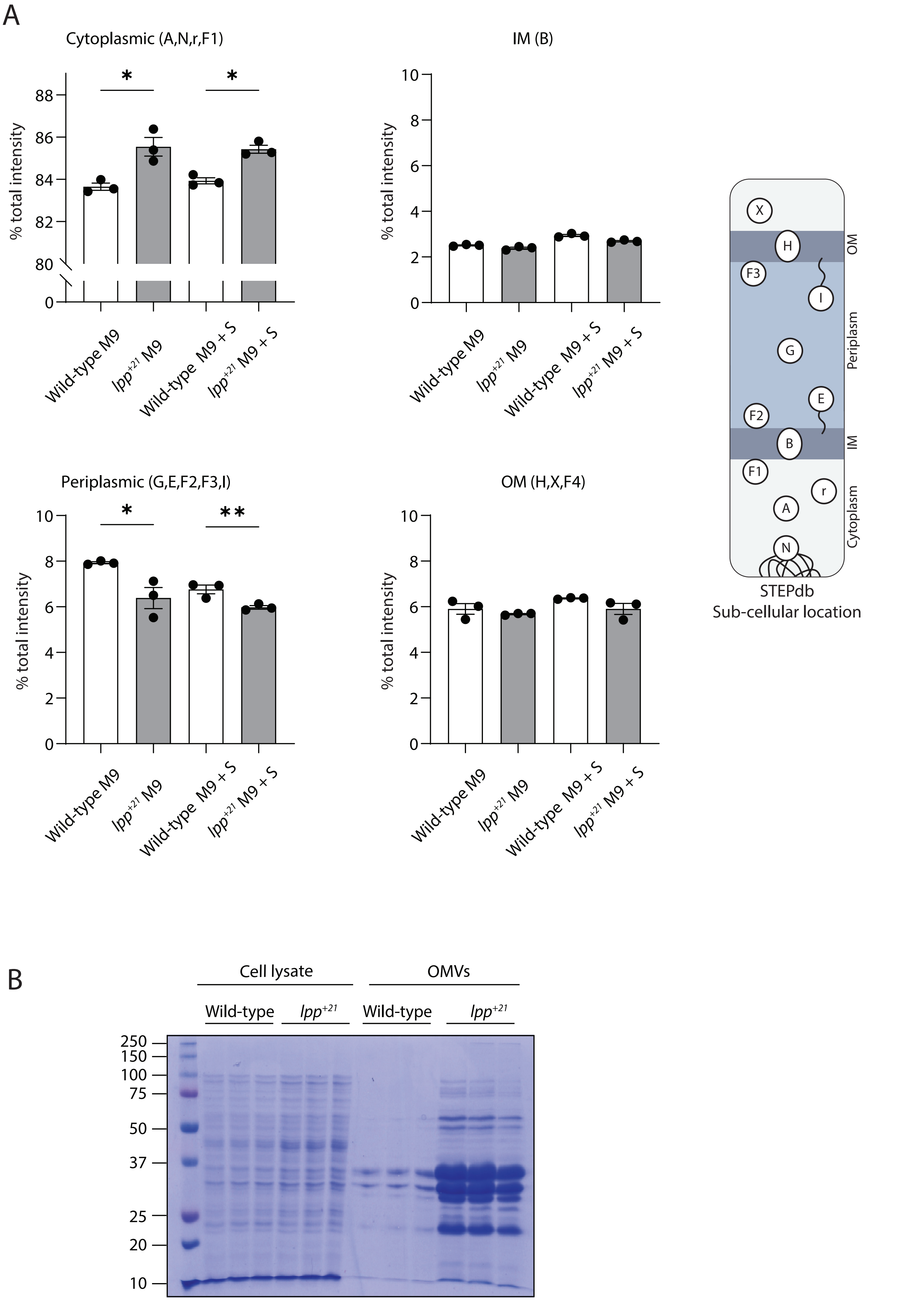
A
Cytoplasmic (A,N,r,F1)
*
*
% total intensity
88
86
84
82
80
0
Wild-type M9
lpp+21 M9
Wild-type M9 + S
lpp+21 M9 + S
IM (B)
10
8
6
4
2
0
Periplasmic (G,E,F2,F3,I)
**
OM (H,X,F4)
OM
Periplasm
IM
Cytoplasm
X
H
F3
I
G
E
F2
B
F1
r
A
N
STEPdb
Sub-cellular location
B
Cell lysate
OMVs
Wild-type
lpp+21
250
150
100
75
50
37
25
20
10

Figure S3

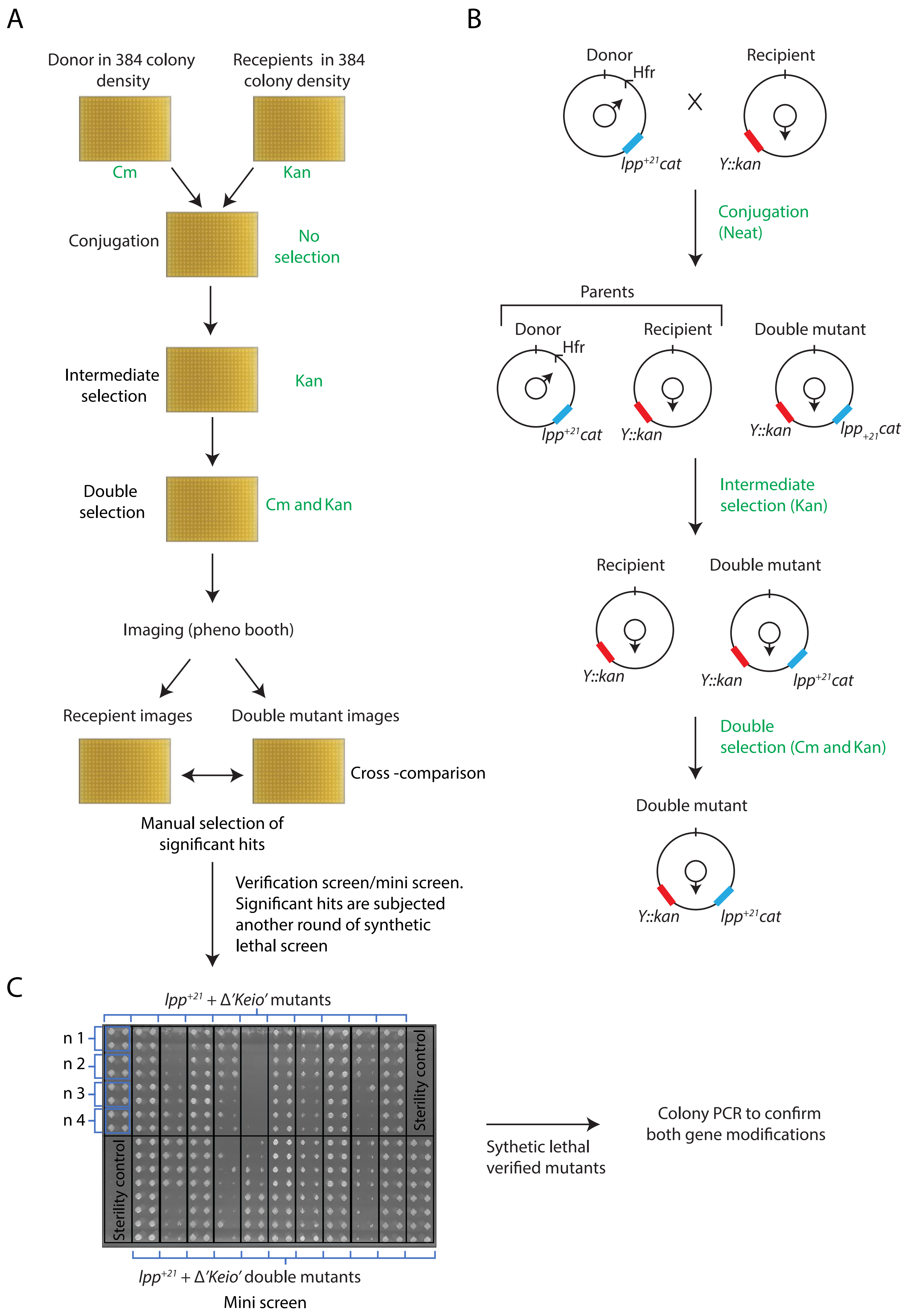
A
Donor in 384 colony density
Recepients in 384 colony density
Cm
Kan
Conjugation
No selection
Intermediate selection
Kan
Double selection
Cm and Kan
Imaging (pheno booth)
Recepient images
Double mutant images
Cross -comparison
Manual selection of significant hits
Verification screen/mini screen. Significant hits are subjected another round of synthetic lethal screen
B
Donor
Hfr
Recipient
lpp+21cat
Y::kan
Conjugation (Neat)
Parents
Donor
Recipient
Double mutant
Intermediate selection (Kan)
Double selection (Cm and Kan)
C
lpp+21 + Δ'Keio' mutants
n 1
n 2
n 3
n 4
Sterility control
lpp+21 + Δ'Keio' double mutants
Mini screen
Sythetic lethal verified mutants
Colony PCR to confirm both gene modifications

Figure S4

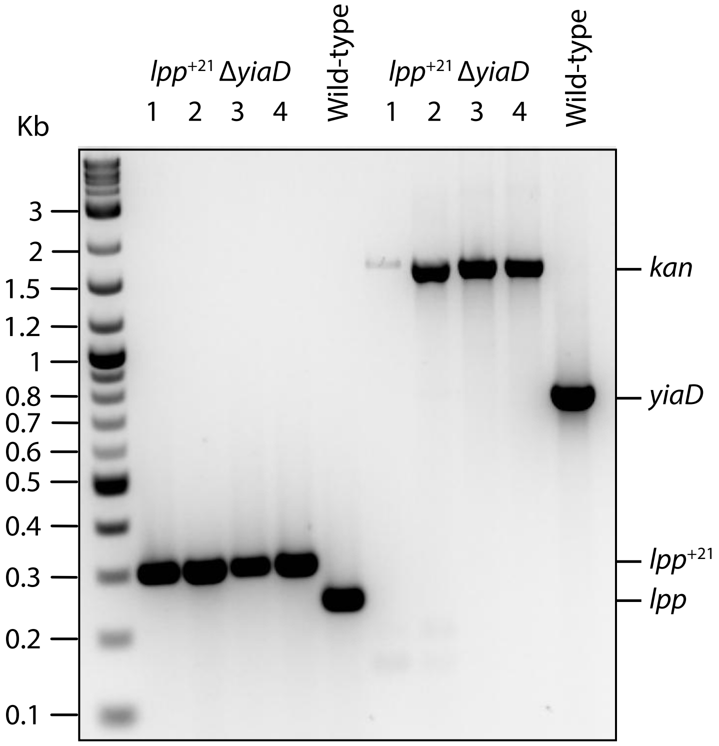
lpp+21 ΔyiaD
1 2 3 4
Wild-type
lpp+21 ΔyiaD
1 2 3 4
Wild-type
Kb
3
2
1.5
1.2
1
0.8
0.7
0.6
0.5
0.4
0.3
0.2
0.1
kan
yiaD
lpp+21
lpp

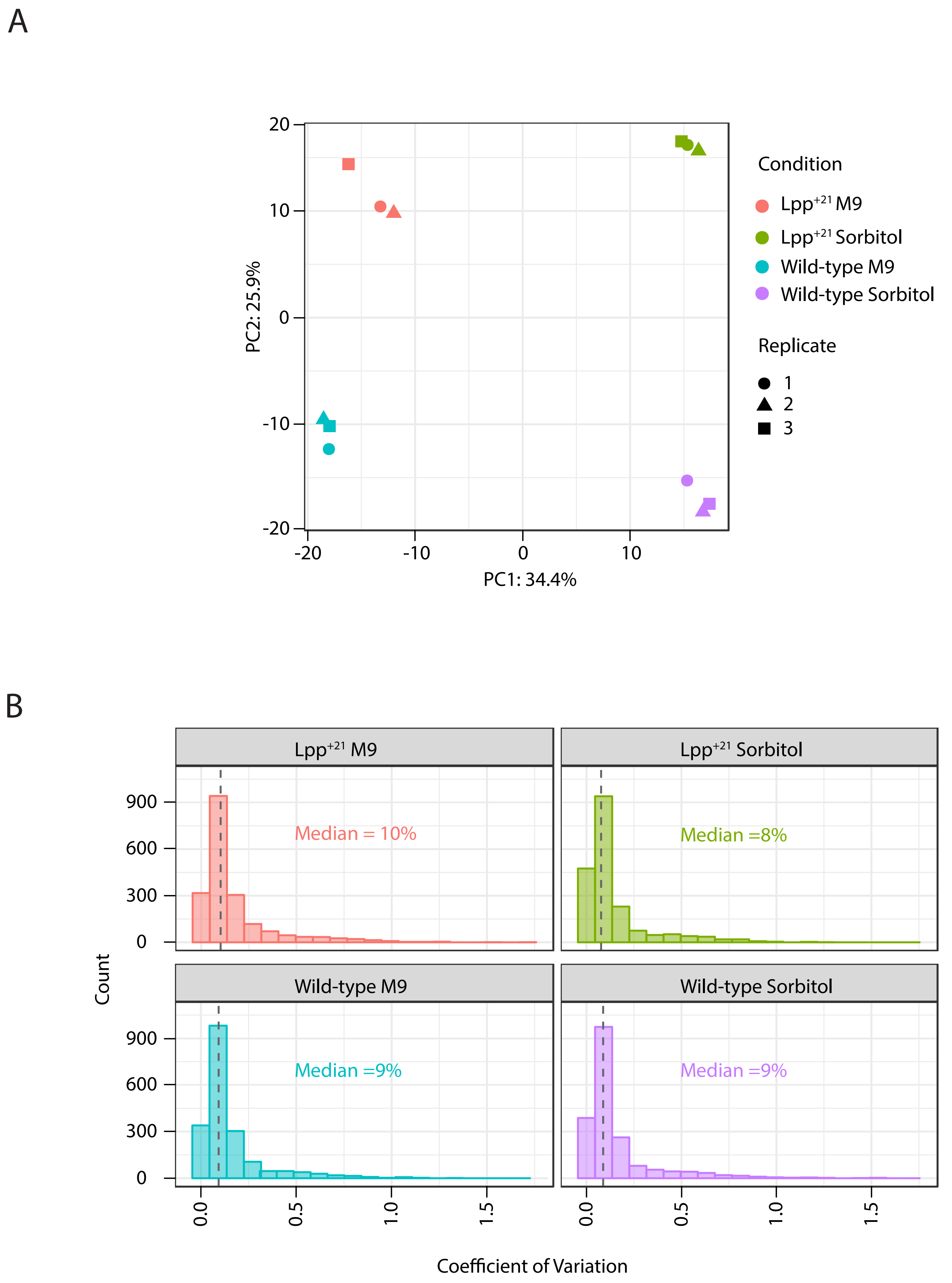
A
Condition
Lpp+21 M9
Lpp+21 Sorbitol
Wild-type M9
Wild-type Sorbitol
Replicate
1
2
3
PC2: 25.9%
PC1: 34.4%
20
10
0
-10
-20
-20
-10
0
10
B
Lpp+21 M9
Lpp+21 Sorbitol
Wild-type M9
Wild-type Sorbitol
Median = 10%
Median =8%
Median =9%
Median =9%
Count
900
600
300
0
0.0
0.5
1.0
1.5
Coefficient of Variation

Figure S6

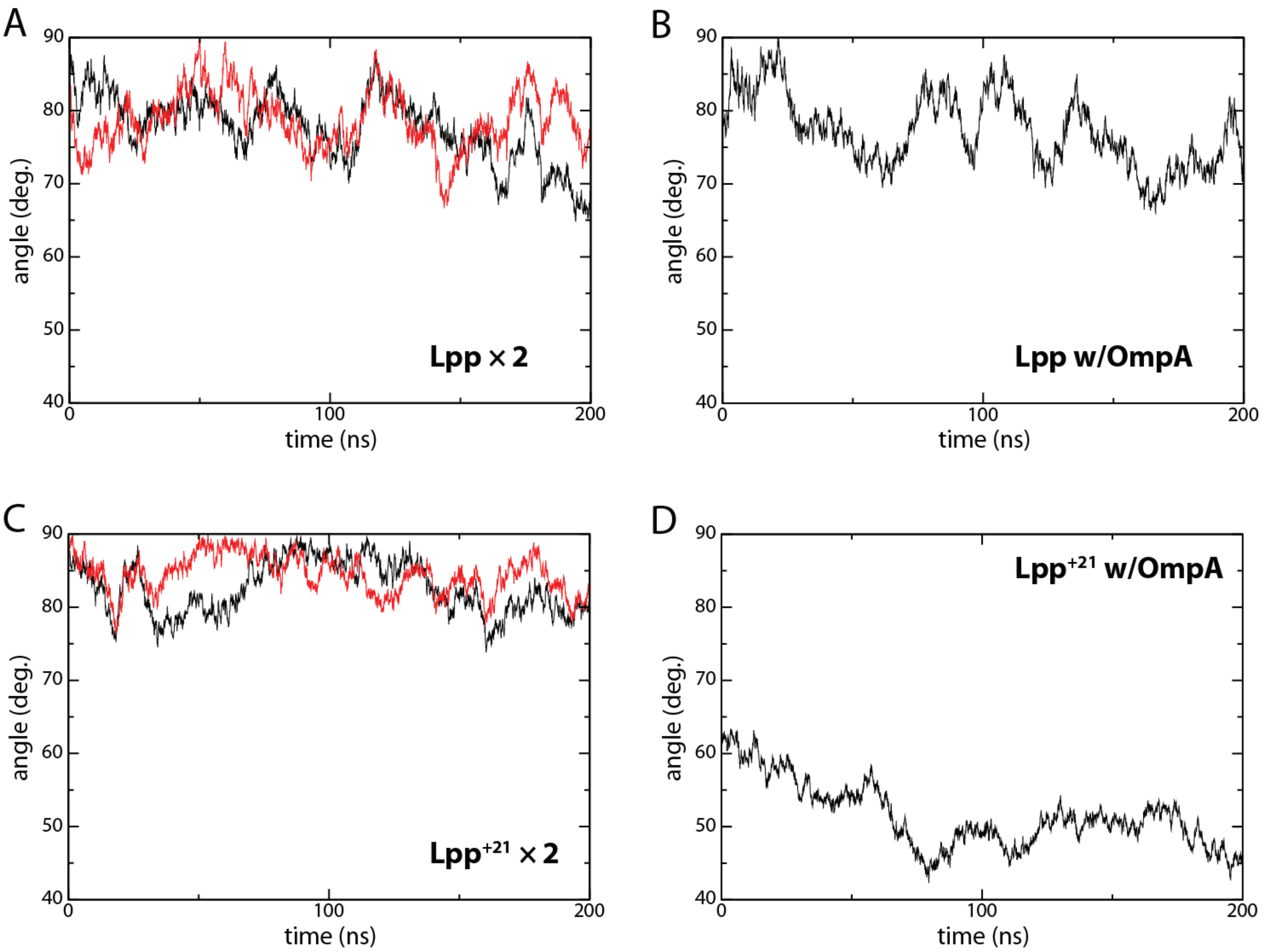

A
B
C
D
Lpp × 2
Lpp w/OmpA
Lpp+21 × 2
Lpp+21 w/OmpA
angle (deg.)
time (ns)
90
80
70
60
50
40
0
100
200


Figure S7
