## Supplemental Table S2 for "Periplasm homeostatic regulation maintains spatial constraints essential for cell envelope processes and cell viability"

**Table S2. Substantive changes in steady-state protein levels in cell envelope of Lpp^+21^**

| **Cellular process** | **Increased** | **Function** |
| --- | --- | --- |
| Peptidoglycan maturation, turnover, remodelling | AmiC | Amidase, cleaves peptide side-chains that link the glycan strands of PG |
| Oligopeptide transport | OppD, OppF, OppC, OppB | PG peptide transmembrane transport (ATP-binding subunit) |
| Ferrous Iron Uptake/transport | FeoAB | Ferrous iron import across the inner membrane |
|  | FhuE, | Feric-coprogen transport, across the OM |
| Stress response | ZraP | Response to cell envelope stress |
|  | YfcG | Response to oxidative stress (has disulphide oxidoreductase activity) |
| IM Protease | RseP/YaeL | Proteolytic cleavage of the anti-sigma factor RseA; Degradation of remnant signal peptides that are left in the inner membrane |
| Protein export | TatA | Twin arginine translocation (Tat) complex |
| c-di-GMP | DgcM/YdaM | c-di-GMP diguanylate cyclase; curli related |
| Other/Unknown | FrdABC | Fumarate reductase complex, in the IM |
|  | DmsB | DMSO reductase, in the IM |
|  | YnfF | Unknown, predicted as a periplasmic protein |
|  | HiuH/YedX | Unknown, predicted as a periplasmic protein |
| **Cellular process** | **Decreased** | **Function** |
| PG-OM bridging | Lpp |  |
| Amino acid transport | DcuA | L-aspartate uptake |
|  | AroP | general aromatic amino acid uptake |
| Carbohydrate transport | GatABCD | Transport and metabolism of Galactitol and DHAP |
|  | SrlE | Transport of sorbitol / mannitol |
|  | MglB | Periplasmic binding protein of a D-galactose/ methyl-D-galactoside ABC transport system |
| Stress response | OsmY | Response to hyperosmotic stress |
| OM integrity | YhdP | Maintaining the outer membrane permeability barrier; interaction with cyclic enterobacterial common antigen |
| AI-2 transport | LsrB | Putative Autoinducer-2 ABC transporter periplasmic binding protein |
| Lipoproteins | BamE | Protein assembly into the outer membrane |
|  | YgdI | Unknown function |
|  | YgeR | Has a LysM domain involved in PG binding; implicated in cell division |
| Lipid | LpxB | lipid A disaccharide synthase, in IM |
|  | GlpQ | Glycerophosphoryl diester phosphodiesterase involved in the utilization of the glycerol moiety of membrane phospholipids |
| Other/Unknown | NuoK | component of NADH dehydrogenase I, in IM |
|  | UbiA | ubiquinone biosynthesis, in IM |
|  | YjiN | Unknown |
|  | YfdI | Unknown |
|  | YfaZ | Unknown, predicted OM Beta barrel |
|  | YahO | Unknown, predicted periplasmic protein |
|  | YdeN | Unknown, predicted periplasmic sulfatase |
|  | YdcS | Unknown, predicted periplasmic binding protein for uncharacterized ABC transporter |
