## Supplemental Table S6 for "Periplasm homeostatic regulation maintains spatial constraints essential for cell envelope processes and cell viability"

**Table S6: Genes up-regulated in $Lpp_{+21}$ strain**

| Gene | Protein | Biological process | $Log_2$ fold change |
|---|---|---|---|
| *lpp/lpp$^{+21}$* | Major outer membrane lipoprotein Lpp | Periplasmic space organisation | 3.03 |
| *insH-1* | CP4-6 prophage; IS5 transposase and trans-activator | transposition of the insertion sequence IS5. | 1.42 |
| *trpD* | anthranilate synthase subunit TrpD | Tryptophan biosynthetic process | 1.20 |

**Method:**

Total RNA was isolated from the cell pellets using an RNAeasy Plus Mini kit (Qiagen) according to the manufacturer's instructions. Extracted RNA samples were quantified using a NanoDrop 100 Spectrophotometer, which was also used to check the quality of the isolated RNA by measuring the sample's A260: A280 ration, expected to be above 1.8. The quality of isolated RNA was also determined by visualisation on agarose gels.

RNA sequencing library was prepared and sequenced at Micromon (Monash University), and the generated fastq files were analysed with RNAsik pipeline (Tsyganov et al., 2018) to produce raw genes count matrix and various quality control metrics, all summarised in MultiQC report (Ewels et al., 2016). For this analysis RNAsik pipeline (Tsyganov et al., 2018) was run with BWA-MEM aligner (Li, 2013) option and reads were quantified with feature counts (Liao et al., 2014). The reference GFF and FASTA files were downloaded from the RefSeq database. The gene counts matrix were then analysed with Degust (Powell, 2019) web tool, including differential expression analysis and several quality plots such as classical multidimensional scaling (MDS) and MA plots. Limma voom (Law et al., 2014) was used in this analysis for model fitting. Degust by Powell, 2019 largely follows limma voom workflow, including the trimmed mean of M values (TMM) normalisation (Robinson & Oshlack, 2010) for RNA composition.
